# Global metabolomic characterizations of *Microcystis* spp. highlights clonal diversity in natural bloom-forming populations and expands metabolite structural diversity

**DOI:** 10.1101/407189

**Authors:** Séverine Le Manach, Charlotte Duval, Arul Marie, Chakib Djediat, Arnaud Catherine, Marc Edery, Cécile Bernard, Benjamin Marie

## Abstract

Cyanobacteria are photosynthetic prokaryotes that are able to synthetize a wild rang of secondary metabolites exhibiting noticeable bioactivity, comprising toxicity. *Microcystis* represents one of the most common cyanobacteria taxa constituting the intensive blooms that arise nowadays in freshwater ecosystems worldwide. They produce numerous cyanotoxins (toxic metabolites), which are potentially harmful to Human health and aquatic organisms. In order to better understand the variations in cyanotoxins production between clones of the *Microcystis*, we investigate the diversity of several strains isolated from the same blooms, from different populations in various geographical area.

Twenty-four clonal strains were compared by genotyping with 16S-ITS fragment sequencing and metabolites chemotyping using LC ESI-qTOF mass spectrometry. While, genotyping can only discriminate between the different species, the global metabolomes reveal clear discriminant molecular profiles between strains. These profiles can be clustered primarily according to their global metabolite content, then to their genotype, and finally to their sampling localities. A global molecular network of all metabolites highlights the production of a wide set of chemically diverse metabolites, comprising only few microcystins, but many aeruginosins, cyanopeptolins and microginins, along with a large set of unknown molecules. They represent the molecular biodiversity that still remain to be investigated and characterized at their structure as well as at their potential bioactivity or toxicity levels.

## 1. Introduction

The frequency and the intensity of cyanobacteria blooms occurring in continental aquatic ecosystems have increased since last decades, due to climate and anthropogenic changes (Carey *et al.*, 2012; Sukenik *et al.*, 2015; Paerl, 2018). These massive cyanobacteria blooms threaten the functioning of aquatic ecosystems through various processes, including an alteration of the trophic network, a decrease of the light penetration within the water column, the decrease of available dissolved oxygen, and also the production of various secondary metabolites potentially toxic for the organisms living in these ecosystems (Carmichael 2008). Indeed, various cyanobacteria genera are able to synthetize a wild range of secondary metabolites (Welker *et al.*, 2012; Shih *et al.*, 2013), with noticeable bioactivity, comprising high toxicity, and which are potentially harmful to Human health and aquatic organisms (Codd et al., 2005; Pearson *et al.*, 2010). These metabolites are also believed to be noticeably involved in the capability of these organisms to proliferate in various environments (Guljamow *et al.*, 2017).

One of the most pervasive bloom-forming cyanobacteria in worldwide freshwater ecosystems is *Microcystis*, as it is now encountered and nowadays proliferate locally in more than 108 countries and on all continents (Sejnohova and Marsalek, 2012; Harke *et al.*, 2016; Ma *et al.*, 2016). Previous documentations also noted that *Microcystis* would occurred during previous decades in only less than 30 countries (Zurawell *et al.*, 2005), suggesting that *Microcystis* is currently proliferating and dominating phytoplankton communities in a wide range of freshwater ecosystems in both temperate and tropical climates. In temperate systems, this organism overwinters in the benthos and during the summer rises to the epilimnion where it can accumulate to form blooms and scums on the water surface (Harke *et al.*, 2016).

Some important features of *Microcystis*, such as buoyancy regulation, storage strategy at the bottom of water column, phosphate (P) and nitrogen (N) uptake capacities and resistance to zooplankton grazing, favour its worldwide spread (Pearl *et al.*, 2011). Indeed, *Microcystis* presents competitive advantages over other cyanobacteria or microalgae in response to nutrient limitation. In addition, many *Microcystis* strains can produce a multitude of bioactive secondary metabolites, including the potent hepatotoxin microcystins (MCs), and thus persistent blooms pose a risk to those who use impaired water resources for drinking water supplies, recreational activities, and fisheries (Codd 2005). Beyond MCs, other toxic compounds or potential secondary metabolites produced by *Microcystis* have also been reported as chemical warfare against grazing of herbivores (Harke *et al.*, 2017).

So far, eleven gene clusters encoding non-ribosomal peptide synthase (NRPS) and/or polyketide synthase (PKS), and two ribosomal ones predicted to be involved in the biosynthesis of secondary metabolites, were found among ten *Microcystis* genomes (Humbert et *al.*, 2013). Seven of these clusters encode enzymes for the biosynthesis of known metabolites (microcystins, aeruginosins, cyanopeptolins, microginins, anabaenopeptins, cyanobactins and microviridins), whereas the six remaining clusters encode enzymes for the biosynthesis of still unidentified products. However, the relationship between cyanobacterial biomass and metabolite concentrations in the environment appears neither systematic nor linear (Briand *et al.* 2002; Liu *et al.*, 2016). Indeed, the production of metabolites, such as microcystins, by *Microcystis* blooms, depends not only on cyanobacterial biomass, but also on the ratio between potentially producing and non-producing genotypes within the population (Via-Ordorika *et al.* 2004).

Despite significant advances in description of the biosynthetic pathways involved in cyanobacterial metabolite production (Dittmann *et al.*, 2015; Wang *et al.*, 2014), the natural functions and the ecological roles played by these molecules are still not well understood (Holland *et al.*, 2013; Zak and Kosakowska, 2016). The biosynthesis of cyanobacterial secondary metabolites consumes a great deal of metabolic energy inducing a significant cost for the cell (Briand *et al.*, 2012). However, natural environment are colonized by various clones exhibiting different corteges of metabolites synthesized (Briand *et al.*, 2009). It has been proposed that the environment it self could favour the selection of *Microcystis* clones that present the metabolite composition that is the more adapted to face the local ecological conditions (Welker *et al.*, 2007; Martins *et al.*, 2009; Agha and Quesada 2014).

Recently, the development of modern approach on mass spectrometry data treatment by molecular networking tools gave us new opportunity for the description of the occurrence and the diversity of cyanobacterial metabolites (Yang *et al.*, 2013; Briand *et al.*, 2016a). With the aim of contributing to better understand the variations in metabolite production, such as microcystins (MCs), between clones of cyanobacterial blooms from different localities, we investigate in the present study the clonal diversity of several *Microcystis* strains isolated from different freshwater bloom-forming populations from various geographical area using such innovative approach based on high resolution mass spectrometry analyses.

## 2. Results

### 2.1. Morphologic and phylogenetic characterization

In order to elucidate the genetic relation between the 24 *Microcystis* strains we have investigated here (Table 1), we performed a first analysis of 1380-pb 16S fragment indicates that all *Microcystis* morpho-species are grouped in a unique and homogenous group, due to high sequence conservation on this fragments (not shown). Using 16S-23S ITS fragments (above 600-bp long), the phylogenetic analysis shows a clear distinction between *M. aeruginosa* and *M. wessenbergii/viridis* morpho-species (Figure 2). Interestingly, the different strains presenting the MC synthesis gene *mcyA* (indicated in red) appear not clustered on the phylogenetic representation, suggesting that be ability of producing MC would be a character disconnected from the phylogeny of strains.

**Table 1:** List of *Microcystis* spp. strains used in this study, their area of origin, the ELISA MC screening, the *mcyA* gene presence and their respective 16S-ITS sequence accession numbers. Stains isolated from the same sample collected the same day from different French area are indicated with: ^a^ = Villerest (2008); ^b^ = Varennes-sur-Seine (2008); ^c^ = Eure et Loire (2010); ^d^ = Valence (2011); ^e^ = Champs – sur-Marne (2012).

| Strain name | Species | Country | Locality/area | MC ELISA<br>detection | *mcyA* PCR<br>detection | 16S-ITS<br>Accession<br>number |
| --- | --- | --- | --- | --- | --- | --- |
| PCC 7806 | *M. aeruginosa* | Netherland | Braakman | + | + | XXXXXX |
| PCC 7820 | *M. aeruginosa* | Scotland | Balgavies | + | + | XXXXXX |
| PMC 95.02<sup>a</sup> | *M. aeruginosa* | France | Villerest | - | - | XXXXXX |
| PMC 98.15<sup>a</sup> | *M. aeruginosa* | France | Villerest | - | - | XXXXXX |
| PMC 155.02 | *M. aeruginosa* | Sénégal | Djoudj | - | - | XXXXXX |
| PMC 156.02 | *M. aeruginosa* | Sénégal | Djoudj | - | - | XXXXXX |
| PMC 241.05 | *M. aeruginosa* | Burkina Faso | Ouahigouya | + | + | XXXXXX |
| PMC 265.06 | *M. aeruginosa* | Burkina Faso | Sian | - | - | XXXXXX |
| PMC 566.08<sup>b</sup> | *M. wesenbergii/viridis* | France | Varennes sur Seine | - | - | XXXXXX |
| PMC 567.08<sup>b</sup> | *M. wesenbergii/viridis* | France | Varennes sur Seine | - | - | XXXXXX |
| PMC 570.08 | *M. aeruginosa* | France | Souppes sur Loing | - | - | XXXXXX |
| PMC 671.10<sup>c</sup> | *M. wesenbergii/viridis* | France | Eure et Loire | - | - | XXXXXX |
| PMC 672.10<sup>c</sup> | *M. wesenbergii/viridis* | France | Eure et Loire | - | - | XXXXXX |
| PMC 673.10<sup>c</sup> | *M. wesenbergii/viridis* | France | Eure et Loire | - | - | XXXXXX |
| PMC 674.10<sup>c</sup> | *M. wesenbergii/viridis* | France | Eure et Loire | - | - | XXXXXX |
| PMC 679.10<sup>c</sup> | *M. aeruginosa* | France | Eure et Loire | + | + | XXXXXX |
| PMC 727.11<sup>d</sup> | *M. aeruginosa* | France | Valence | - | - | XXXXXX |
| PMC 728.11<sup>d</sup> | *M. aeruginosa* | France | Valence | + | + | XXXXXX |
| PMC 729.11<sup>d</sup> | *M. aeruginosa* | France | Valence | + | + | XXXXXX |
| PMC 730.11<sup>d</sup> | *M. aeruginosa* | France | Valence | - | - | XXXXXX |
| PMC 807.12<sup>e</sup> | *M. wesenbergii/viridis* | France | Champs sur Marne | + | + | XXXXXX |
| PMC 810.12<sup>e</sup> | *M. aeruginosa* | France | Champs sur Marne | - | - | XXXXXX |
| PMC 816.12<sup>e</sup> | *M. aeruginosa* | France | Champs sur Marne | + | + | XXXXXX |
| PMC 826.12<sup>e</sup> | *M. aeruginosa* | France | Champs sur Marne | - | - | XXXXXX |

**Figure 1:**
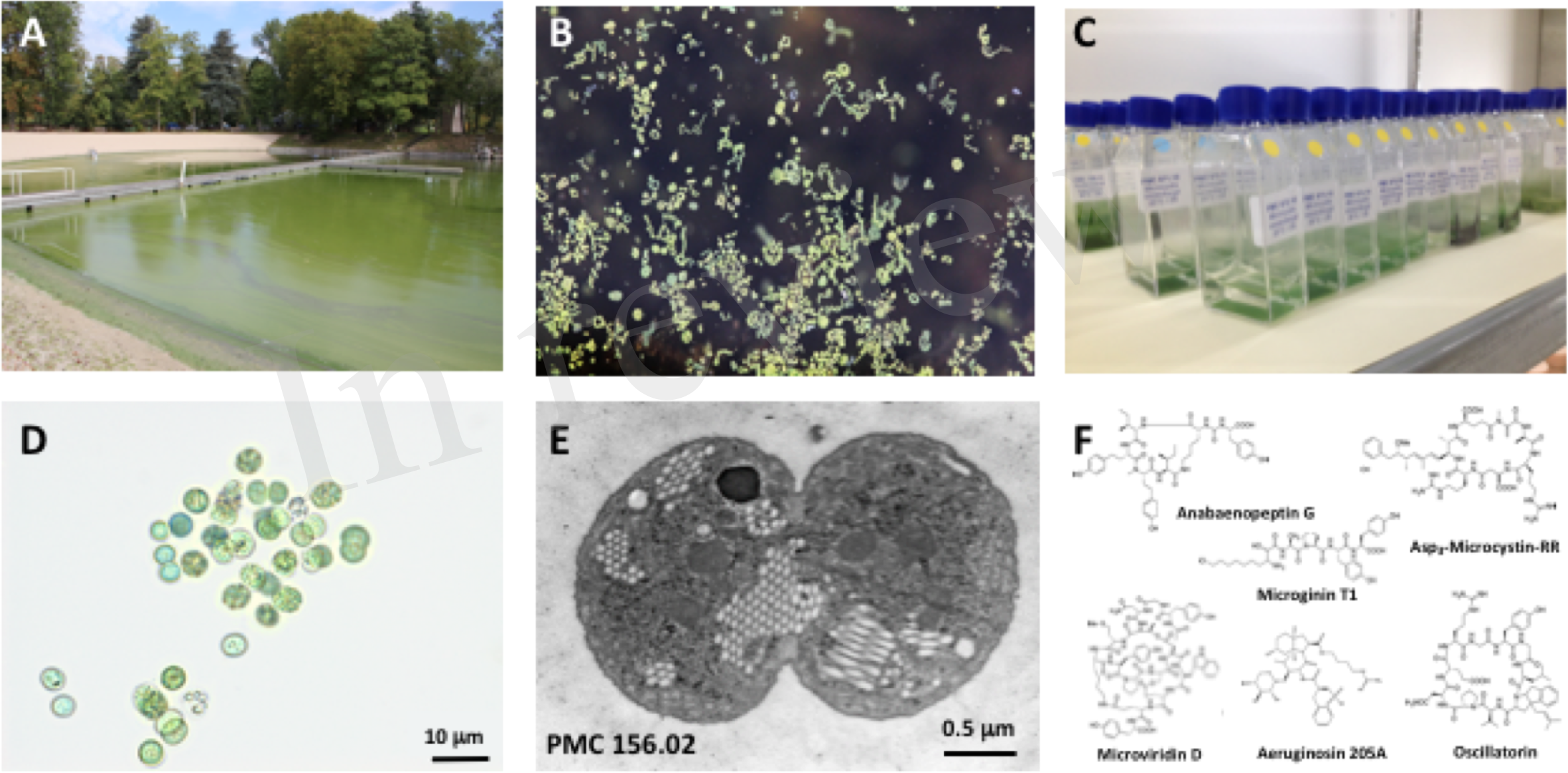
Microcystis *spp*. General view of a representative intense *Microcystis* sp. bloom in a recreational pound (Champs-sur-Marne, © B. Marie) (A). Macrograph of *Microcystis* colonies at surface water (© B. Marie) (B). Example of 15-mL vessels containing the monoclonal strains of *Microcystis* spp. maintained in the Paris’ Museum Collection (PMC) of cyanobacteria (MNHN, Paris, © C. Duval) (C). Example of micrograph of the isolated monoclonal culture of the *Microcystis aeruginosa*, where scale bare represents 10 μm (© C. Duval) (D). Representative picture of *Microcystis aeruginosa* cell from PMC 156.02 strain (here in division) under transmission electron microscope, where scale bare represents 0.5 μm (© C. Djediat) (E). General structures of various cyanobacterial metabolites belonging to the microcystin, anabaenopeptin, microginin microviridin, aeruginosin and oscillatorin families (F).

**Figure 2:**
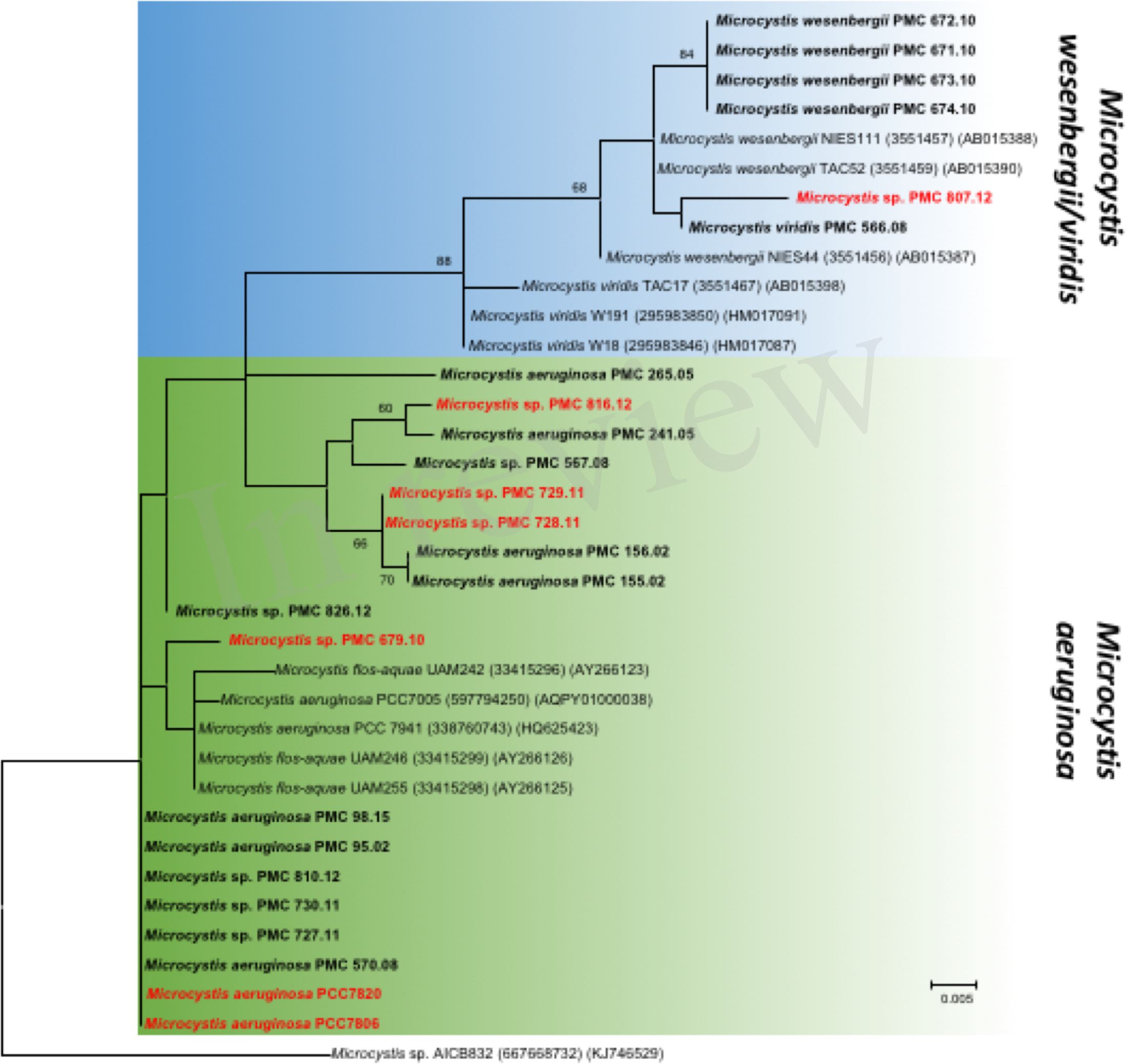
Maximum likelihood (ML) tree based on partial 16S-23S ITS sequences. The sequences obtained in this study are indicated in bold. The strains, which exhibit *mcyA* gene, are indicated in red. Other sequences were retrieved from GenBank, accession numbers in brackets. Bootstrap values >60% are shown at the nodes. The scale bar indicates number of nucleotide substitutions per site. The *Microcystis* sp. AICB832 was used as an out-group.

### 2.2. Global metabolome analyses

The metabolomic shotgun analyses reveal discriminant metabolic profiles between strains collected from both different or identical sites. Whereas previous works had highlighted the metabolic diversity of some *Microcystis* strains based on few identified metabolites or cyanotoxines (Martins *et al.*, 2009; Welker *et al.*, 2004; Welker *et al.*, 2006), we present here a global picture of the metabolome of each strain. Using HR ESI-Qq-TOF with the 24 *Microcystis* strain extracts, 2051 distinct mass ions in a range of 400-2000 Da were recorded (*i.e.* with a signal to noise ratio in excess of 6, and respective relative peak intensity superior to 5000-count in at least one sample threshold), each strain exhibiting between above 100 and 300 different main metabolites of reliable (>10 000 counts) respective intensity (Figure 3; supplementary figure S1). A hierarchical clustering was performed according to Bray-Curtis indexes calculated between all 24 strains according to the relative intensity of each analytes in each strains. This representation (Figure 3) clearly shows a clustering of all strains producing MCs (*mcyA*+/MC+, in red), on one side, and of other strains not producing MCs (*mcyA*−/MC−, in grey). This clustering shows that some strains from the same environment exhibits very similar metabolite fingerprints (*e.g.* PMC 728.11 and 729.11), when other from the same localities exhibits much more dissimilar metabolite fingerprints (*e.g.* PMC 728.11 and 727.11), being more similar with strains from faraway locations (*e.g.* PMC 729.11 and 816.112). Additional non-metric multidimensional scaling (nMDS) and PERMANOVA analyses based on Bray-Curtis index indicates that the ability to produce MC seems to be the first main driver of the global metabolome of *Microcystis* molecular fingerprinting, the species and the localities representing less explicative parameters (supplementary figure 2).

**Figure 3:**
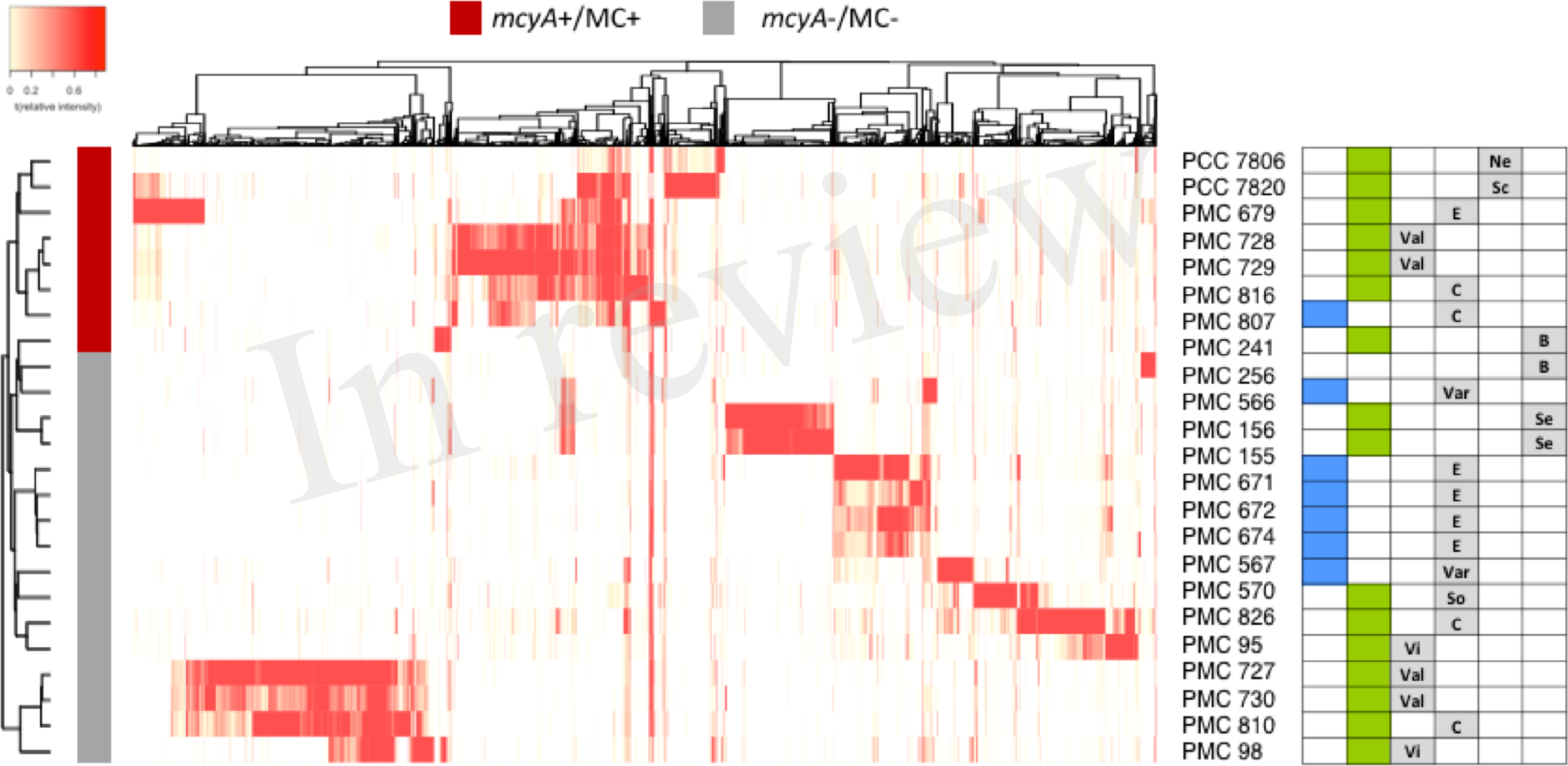
Heatmap representation of the metabolome of the 24 *Microcystis* spp. monoclonal strains analysed using HR ESI-TOF, representing 2051 different analytes (present in at least three strains, with minimal peak intensity > 5000 counts) identified by MetaboScape software. The hierarchical clustering between strains was performed according to Bray-Curtis distance method. Blue and green squares indicate *M. aeruginosa* and *M. wesenbergii/viridis*, respectively. Sampling localities are: C=Champs-sur-Marne; B=Burkina Faso; E=Eure et Loire; Ne=Netherlands; Sc=Scotland; Se=Senegal; So=Souppes-sur-Loin; Var=Varennes-sur-Seine; Val=Valence; Vi=Villerest.

### 2.3. Metabolite molecular network

A molecular network based on the global fragmentation pattern profile of all observed metabolites in the 24 strains investigated. The GNPS algorithm automatically compares all MS/MS spectra by aligning them one by one, grouping identical molecules (presenting identical mass and fragmentation pattern) and assigning cosine score ranking from 0 to 1 to each alignment, allowing network reconstruction of the link between each molecule according to the cosine score links between all molecules with a cosine score significance threshold set to 0.6. The resulting network constituted of a total of 925 nodes from the 1374 different analytes which MS/MS data have been obtained (supplementary figure 1), represents a starting point for the annotation of unidentified metabolites, according to respectively identified molecules within the different clusters, and to their occurrence in *M. aeruginosa* and/or *M. wesenbergii/viridis* strains (Figure 4).

**Figure 4:**
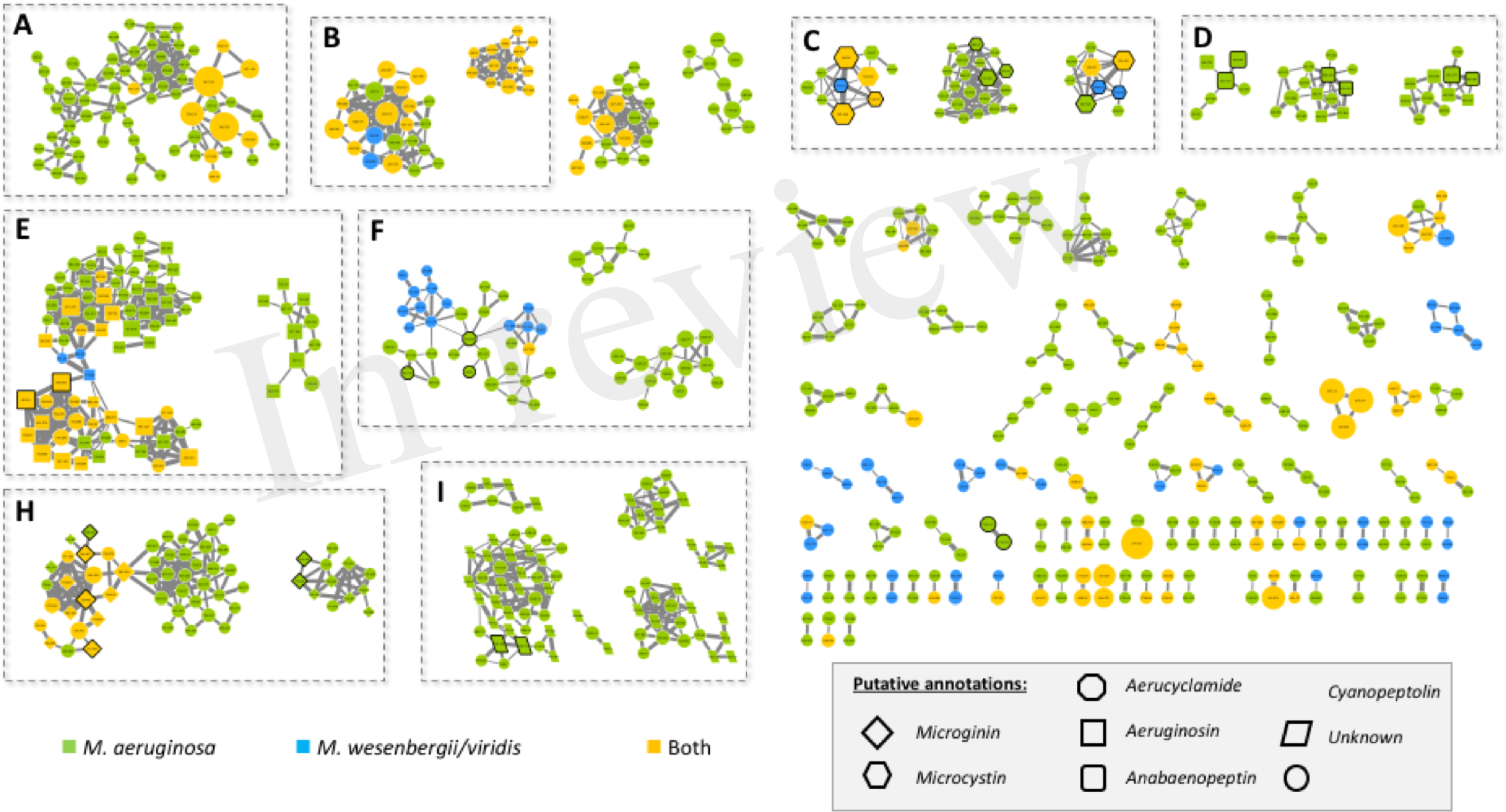
Molecular network generated from MS/MS spectra from the 24 *Microcystis* strains using GNPS tool (all data and results are freely available on the GNPS server at the address http://gnps.ucsd.edu/ProteoSAFe/status.jsp?task=c017414365e84334b38ae75728715552). The nodes of the analytes detected in *M. aruginosa* or *M. wessenbergii/viridis* strains only are indicated in green and blue, respectively, when analytes detected in both species are indicated in orange. Uncharacterized analytes are indicated by circles constituting potential new analogues. Analytes whom individual masses match with known secondary metabotites from cyanobacteria (listed in supplementary table 1) are indicated as specific shapes. Standard molecules analyses similarly are indicated by a heavy black perimeter. Only cluster regrouping at least 2 analytes are represented.

The *Microcystis* strains produce a wide set of chemically diverse metabolites, which principal clusters can attemptedly been identified thanks to analytical standards available for some cyanobacterial secondary metabolite family, or to match with components from libraries publically available from GNPS platform, such as HMDB, NIST14 or METLIN. Considering the strains producing these analytes, and above a quarter of them appears to be specific of *M. wessenbergii/viridis* strains, when more than half are specific of *M. aeruginosa*, the rest being observed on both species.

The grouping of different analytes in the same molecular cluster is based on similarity of their fragmentation patterns, each cluster being potentially specific of the structure of chemical families. Among those larger clusters, we were thus able to annotate some of them, being constituted by ions of small metabolites, such as di-peptides and small peptides (1 cluster in A area), of microcystins (3 clusters in C area), of anabaenopeptins (3 clusters in D area), of aeruginosins (2 clusters in E area), of aerucyclamides (3 clusters in F area), of microginins (2 clusters in H area) and cyanopeptolons (6 clusters in I area), together with various clusters of unknown components, comprising non-identified ions (for example 2 clusters in B area). Less than a third of the metabolites observed here could be annotated, thanks to their respective mass and fragmentation patterns when compare to those of the above 800 metabolite of freshwater cyanobacteria described so far and listed in supplementary table 1. These unidentified ions that belong to annotated clusters are then considered as potential new analogues of their respective molecular family.

### 2.4. Known cyanobacteria secondary metabolite clusters

#### Microcystins

Microcystins are cyclic heptapeptides that have been firstly described from *M. aeruginosa*. Above 250 different variants have been described so far (Catherine *et al.*, 2017), 138 being references in our database for cyanobacterial metabolite (supplementary table 1). They are characterized by the presence of a non-proteinacous amino acid in position 5 (Adda), two amino acid derived from Asp and Glu in position 3 and 6, respectively, and 2 very variable positions (2 and 4), that serve as reference to the name of the variant. Three microcystis clusters were highlighted according to the presence of 5 standard molecules (Dmet(Asp3)-MC-LR, MC-LR, MC-YR, MC-LA and MC-HtyR) analyses in parallel of the 24 *Microcystis* extracts with the same protocol (Figure 5). Other components of these clusters correspond to ions presenting a match of their respective mass with those of other MC variants previously described (supplementary table 1), or for ¼ of them to potential new analogues. Observation of their respective MS/MS spectra shows that they present distinct but similar fragmentation patterns of other known MC variants.

**Figure 5:**
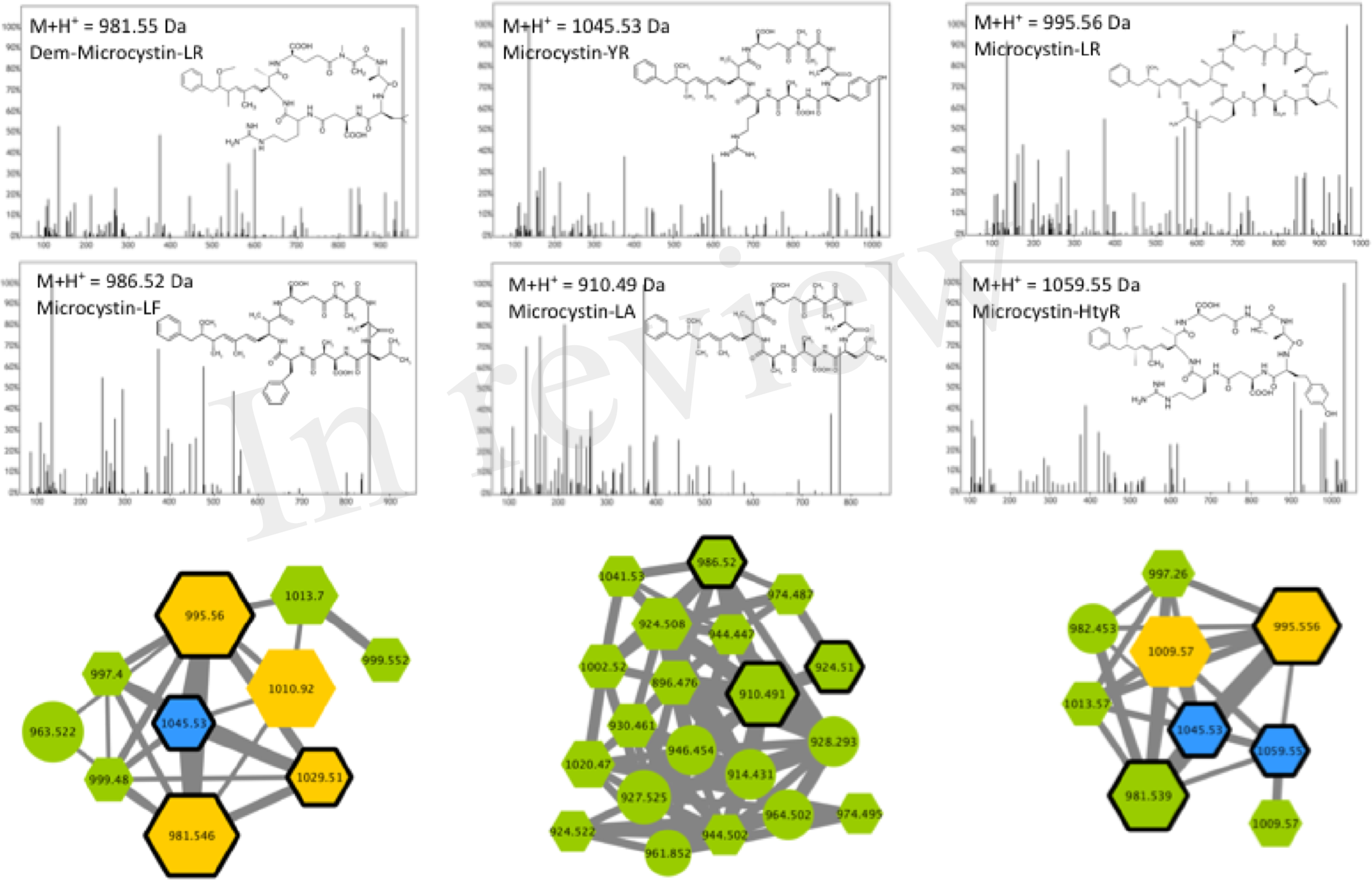
Microcystins clusters highlighted by the GNPS analysis based on the MS/MS CID fragmentation spectra obtained from the 24 *Microcystis* strains. Analytes detected in *M. aeruginosa* or *M. wessenbergii/viridis* strains only are indicated in green and blue, respectively, when analytes detected in both genera are in orange. Analytes whom individual masses match with known microcystins are indicated as hexagons. Standard molecules analyses similarly are indicated by a heavy black perimeter. Example of MS/MS spectra and chemical structures are shown for (Asp3)-microcystin-LR, microcystin-YR, -LR, -LF, -LA and -HtyR. Notice that (M+H)^+^ and (M+2H)^2+^ ions may be grouped in distinct clusters.

#### Aeruginosins

Aeruginosins constitute a linear tetrapeptide family that have been firstly described from *M. aeruginosa*, and that represent above 94 different variants that have been described so far (Supplementary table 1). Their MS/MS fragmentation patterns are often characterized by the presence of a Choi fragment (immonium with 140.109 *m/z*) and other recurrent fragments from Hpla or Pla. Their composition is rather variable and the component of this family exhibit masses comprised between 430 and 900 Da (Welker and von Döhren 2006). The molecular network obtained from the 24 *Microcystis* strains exhibits 2 aeruginosin clusters (Figure 6) that were highlighted by to the presence of 2 standard molecules (aeruginosin 98A and 98B). Other components of these clusters correspond to ions presenting a mass match with other variants of aeruginosin previously described (supplementary table 1), or for ½ of all this compounds to potential new analogues, that aim at being characterized now, in further dedicated works.

**Figure 6:**
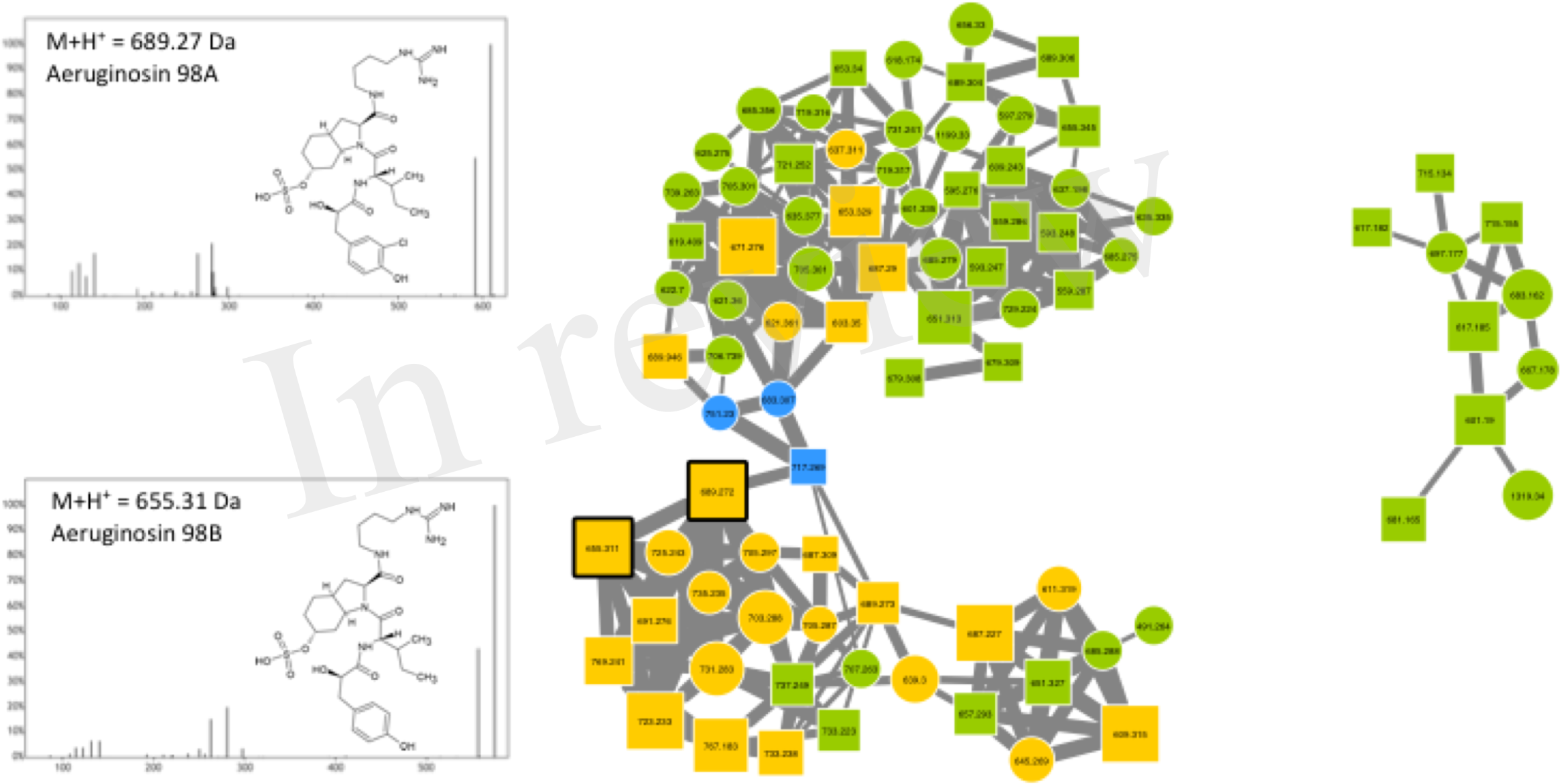
Aeruginosin cluster highlighted by the GNPS analysis based on the MS/MS CID fragmentation spectra obtained from the 24 *Microcystis* strains. Analytes detected in *M. aeruginosa* or *M. wessenbergii/viridis* strains only are indicated in green and blue, respectively, when analytes detected in both genera are in orange. Analytes whom individual masses match with known aeruginosins are indicated as scares with right corners. Standard molecules analyses similarly are indicated by a heavy black perimeter. Example of MS/MS spectra and chemical structures are shown for aeruginosin 98A and 98B.

#### Anabaenopeptins

Anabaenopeptins constitutes a very diverse family of cyclic hexapeptides that have been described from now from *Microcystis*, *Planktothrix*, *Anabaena*, *Aphanizomenon* and *Nostoc*. Above 75 different variants have been described so far (supplementary table 1). They are characterized by the presence of a peptide bound between the D-Lys placed in position 2, and the carboxylic group of the amino acid placed in position 6. Except fro the D-Lys (position 2) all other positions are variable allowing a large structural diversity of the family which molecules exhibit masses between 750 and 950 Da (Welker and von Döhren 2006). Three anabaenopeptin clusters were highlighted here (Figure 7) according to the presence of 4 standard molecules (anabaenopeptin A, B, E and oscyllamide Y). Other components of these clusters correspond to ions presenting a mass with other variants previously described (supplementary table 1), or for ½ of all of them to compounds that very likely correspond to potentially new analogues. All observed anabaenopeptin compounds are from *M. aeruginosa* strains suggesting that *M. wessenbergii/viridis* strains are not capable of the synthesis of molecules of this family and may not possess the corresponding *apt* synthetic gene cluster.

**Figure 7:**
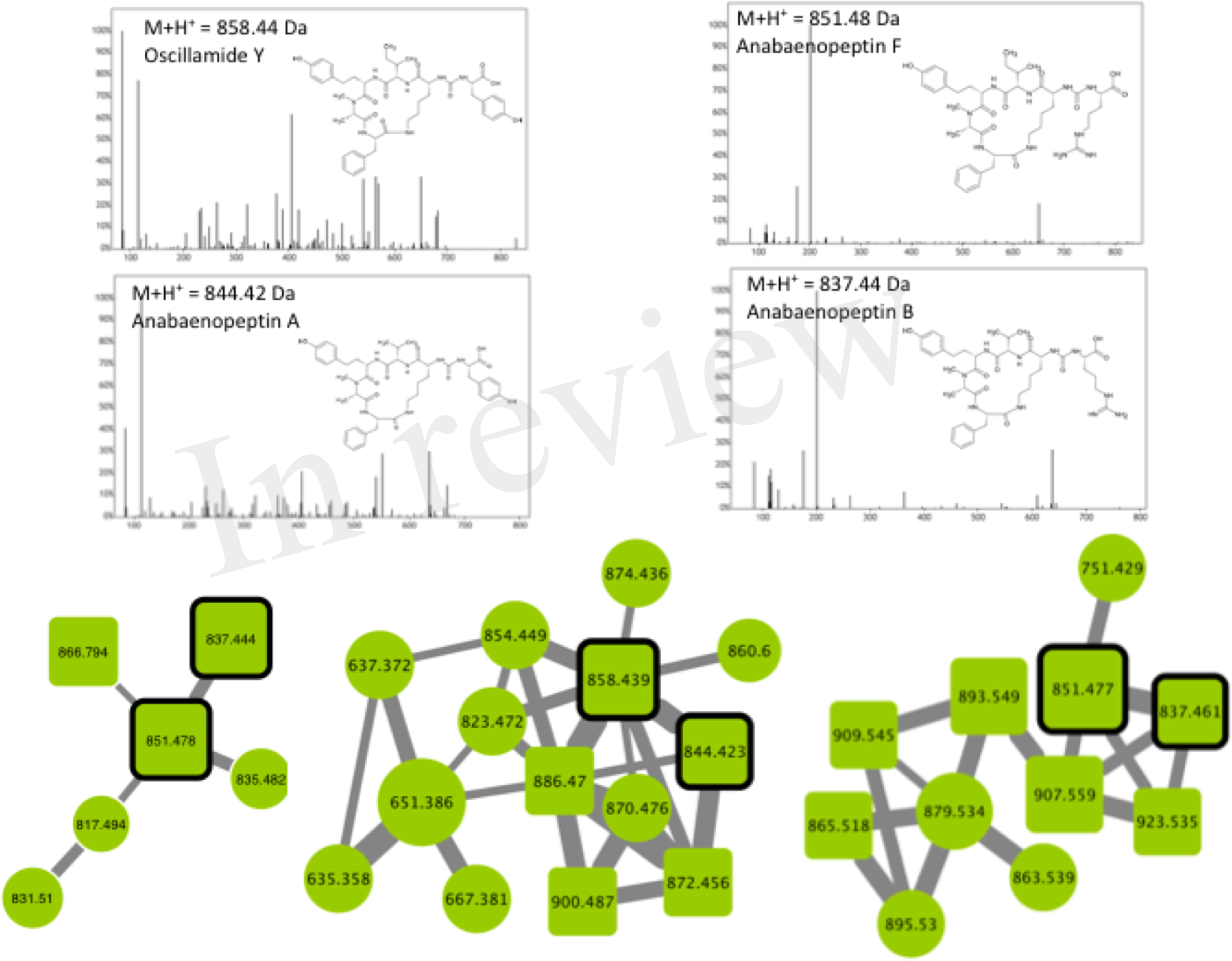
Anabaenopeptin clusters highlighted by the GNPS analysis based on the MS/MS CID fragmentation spectra obtained from the 24 *Microcystis* strains. Analytes detected in *M. aeruginosa* strains are indicated in green. Analytes whom individual masses match with known anabaenopeptin are indicated as scares with round corners. Standard molecules are indicated by a heavy black perimeter. Uncharacterized analytes are indicated by circles constituting potential new analogues. Example of MS/MS spectra and chemical structures are shown for anabaenopeptin A, B, F and Oscillamide Y. Notice that (M+H)^+^ and (M+2H)^2+^ ions may be represented by different nodes grouped in distinct clusters.

#### Cyanopeptolins

Cyanopeptolins belong to a large family of cyclic depsipeptides, that also comprises micropeptins and aeruginopeptins, representing above 170 variants. Those molecules are characterized by the presence of the non-proteinaceous amino acid Ahp and by a six-aa long ring formed by an ester bound between Thr or Pro in position 1 and the carboxylic group of the N-terminal amino acid (position 6). The lateral chain can exhibit a variable length and is constituted by one or two amino acid and potentially linked to an aliphatic fatty acid (Welker and von Döhren 2006). In the molecular network, 2 analytes of the cyanopeptolin clusters correspond to 2 standard molecules (cyanopeptolin B and D), and various other components correspond to ions presenting a mass that corresponds to different variants previously described (supplementary table 1), allowing us to annotate them as cyanopeptolin specific clusters (Figure 8). Above ½ of the analytes present in these clusters correspond to unknown compounds constituting potential new analogues. We observe here that all these cyanopeptolin compounds are from *M. aeruginosa* strains suggesting that *M. wessenbergii/viridis* strains are not capable of the synthesis of molecules of this family and may not possess the corresponding *mcn/oci* synthetic gene cluster.

**Figure 8:**
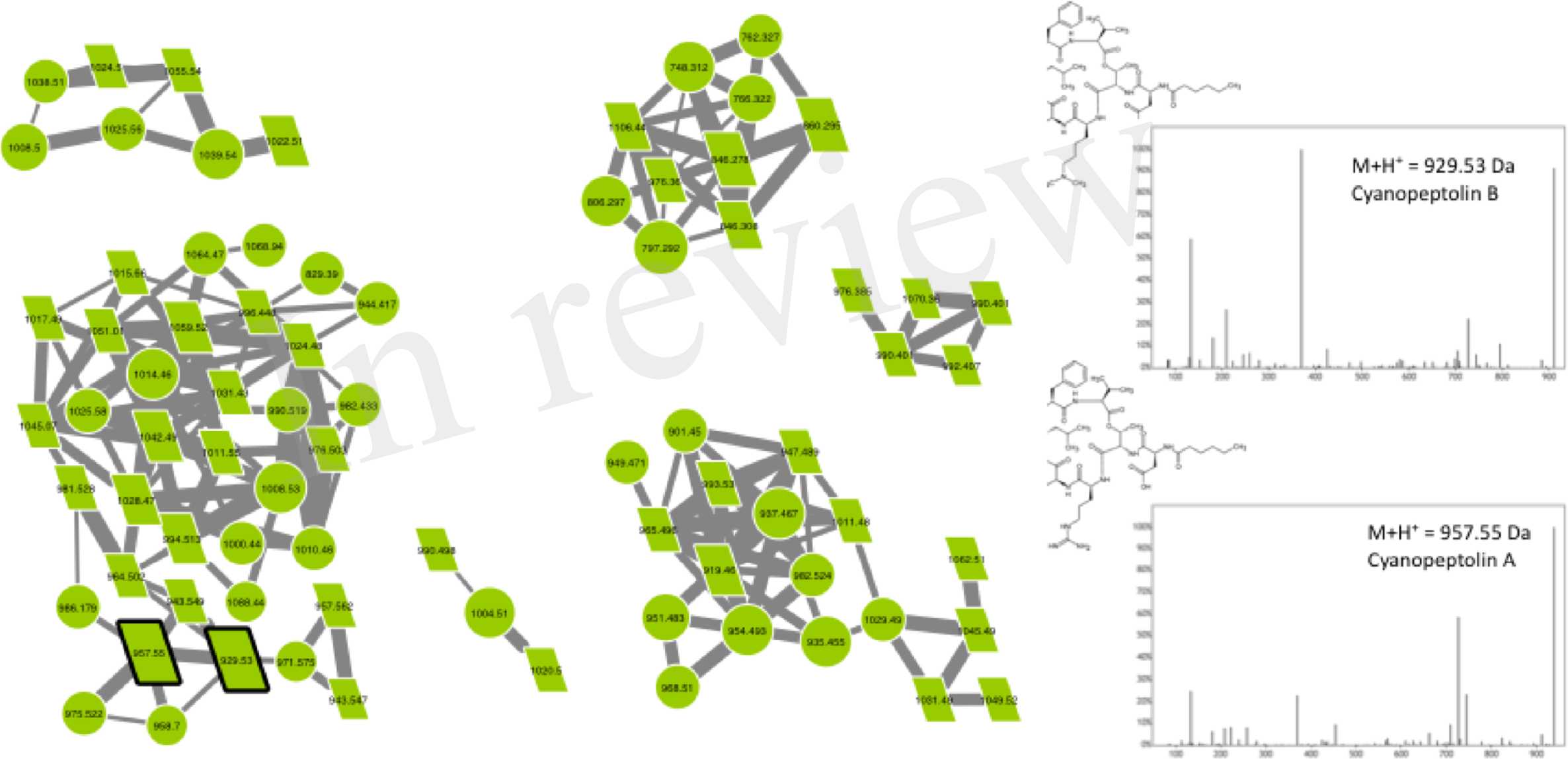
Cyanopeptolin clusters highlighted by the GNPS analysis based on the MS/MS CID fragmentation spectra obtained from the 24 *Microcystis* strains. Analytes detected in *M. aeruginosa* strains are indicated in green. Analytes whom individual masses match with known cyanopeptolin are indicated as parallelepiped. Standard molecules are indicated by a heavy black perimeter. Uncharacterized analytes analyses similarly are indicated by circles constituting potential new analogues. Example of MS/MS spectra and chemical structures are shown for cyanopeptolin A and B.

#### Microginins

Microginins are linear pentapeptides (the length of the sequence varying from 4 to 6 amino acids) initially identified from *Microcystis aeruginosa*, then from other species and in other genera such as *Planktothrix* (Welker and von Döhren 2006). These molecules are composed by a characteristic non-proteinaceous amino acid Ahda in N-terminy, the other position bearing variable amino acid structures, comprising Tyr, Pro Hty, Trp, Ala, Ser…. Relatively few microginin variants (less than 40) have been described so far (supplementary table 1). According to the molecular network, above two third of the analytes present in the two microginin clusters correspond to unknown compounds constituting potential new analogues, when six standard molecules could have been retrieved in our analysis (microginin 757, 711, BN578, FR1, FR2 and SD755).

**Figure 9:**
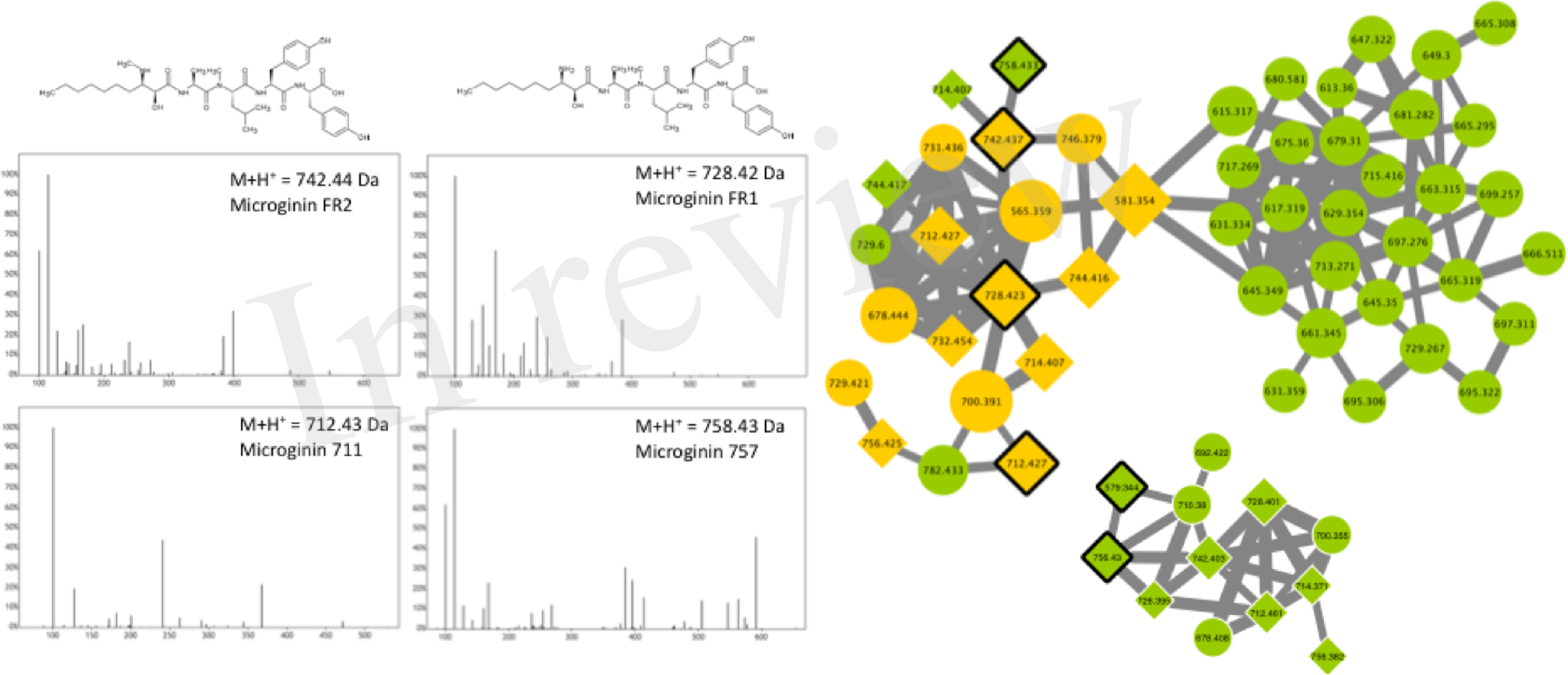
Microginin clusters highlighted by the GNPS analysis based on the MS/MS CID fragmentation spectra obtained from the 24 *Microcystis* strains. Analytes detected in *M. aeruginosa* strains only are indicated in green and blue, when analytes detected in both genera are in orange. Analytes whom individual masses match with known microginin are indicated as 45°-tilted scares. Standard molecules analyses similarly are indicated by a heavy black perimeter. Uncharacterized analytes are indicated by circles constituting potential new analogues. Example of MS/MS spectra and chemical structures are shown for microginin FR1, FR2, 711 and 757.

#### Aerucyclamides/cyanobactins

The name “cyanobactin” has been proposed in order to grouped all cyclic peptides containing proteinogenous amino acids that are post-translationally modified in heterocyclic amino acids and isoprenoid derivatives (Sivonen *et al.*, 2010). It comprises various cyclamides (cyclic peptides of 6 amino acids) that have been identified in freshwater cyanobacteria such as *Microcystis*, *Planktothrix* and *Nostoc*, but also in symbiotic cyanobacteria species. More than 30 variants have been described so far, but more molecules could be related to the family that represent a very large variety of chemical structures (Martins and Vasconcelos 2015). Three different cyanobactin clusters were observed in the molecular network, comprising three aerucyclamide standard molecules (aerucyclamide A, B and C) that were identified by GNPS tool. Above ¾ of the analytes from these clusters representing potential new analogues, that need to be characterized by further dedicated analyses.

**Figure 10:**
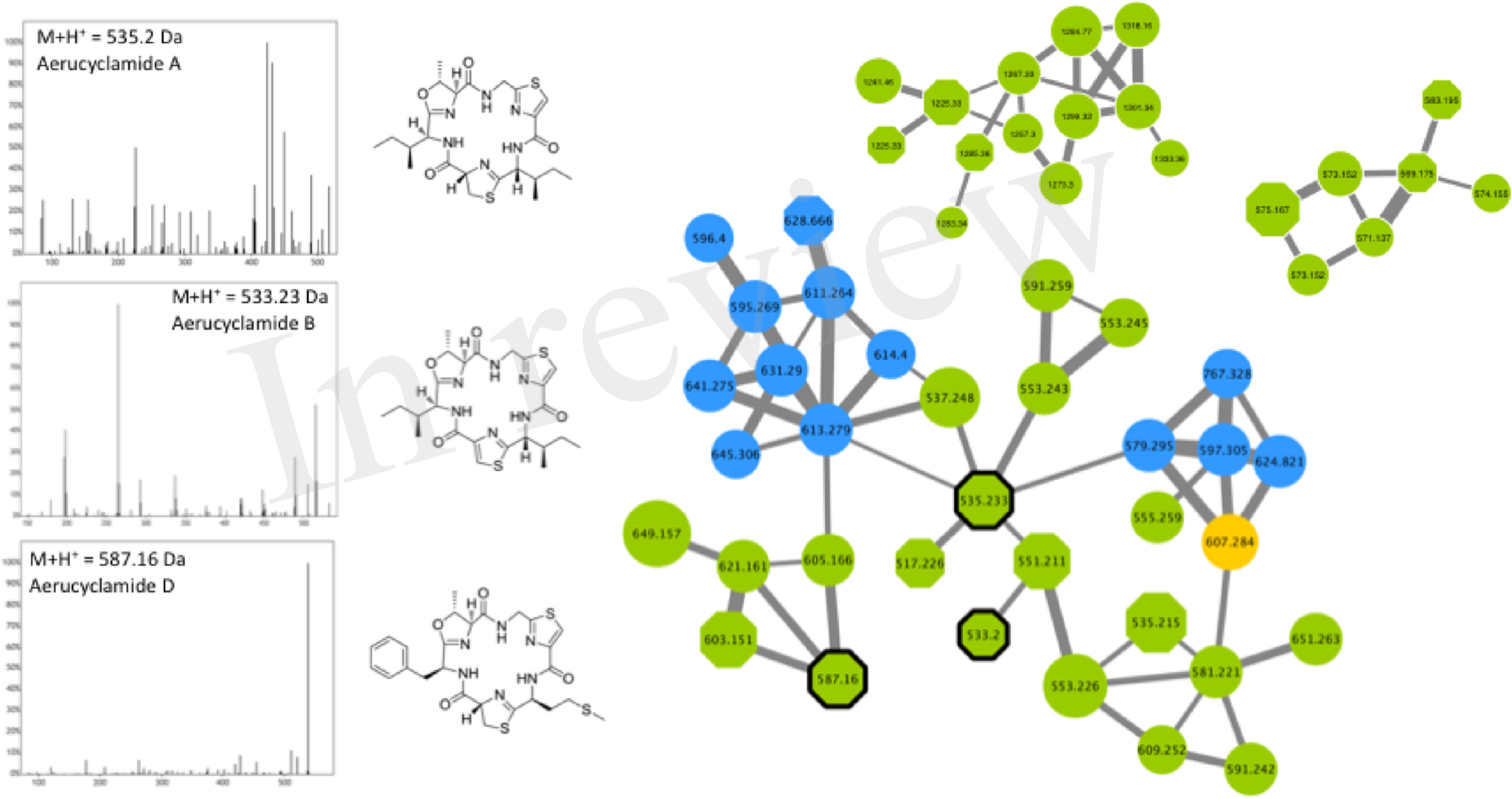
Aeruclyclamide clusters highlighted by the GNPS analysis based on the MS/MS CID fragmentation spectra obtained from the 24 *Microcystis* strains. Analytes detected in *M. aeruginosa* or *M. wessenbergii/viridis* strains only are indicated in green and blue, respectively, when analytes detected in both genera are in orange. Analytes whom individual masses match with known aerucyclamides are indicated as octogons. Uncharacterized analytes are indicated by circles constituting potential new analogues. Standard molecules analyses similarly are indicated by a heavy black perimeter. Example of MS/MS spectra and chemical structures are shown for aerucyclamide A, B and D.

### 2.5. Uncharacterized cyanobacteria metabolite clusters

#### Potential primary metabolite clusters?

Various other important clusters comprising few tens of analytes, that are mostly present in high amount (higher peak intensity showed by larger circles) in a large set of strains from both *M. aeruginosa* and *M. wessenbergii/viridis* (Figure 4). These compounds present molecular masses comprised between 300 and 500 Da, suggesting that it could correspond to relatively small compounds (supplementary figure 3). We can speculate, that these components could correspond to primary metabolites used for the general metabolism of various strains, however additional effort should be made in order to propose annotations for these ubiquitous molecules present in these specific clusters. Interestingly, the compounds from these latter cluster present high fragmentation patterns, illustrated with various similarity and high cosine score (numerous and tick links between nodes), suggesting they should present very similar structures.

#### Uncharacterized secondary metabolite clusters?

Interestingly, other clusters correspond to unknown components, some of them being only synthesis by *M. wessenbergii/viridis* as exemplified in supplementary figures 4 and 5. These compounds which molecular mass are relatively high (between 950 and 1300 Da) might correspond to specific secondary metabolite families, belonging to other known family of cyanobacterial metabolite, but poorly characterized, such as microviridins and/or aeruginoguanidins, for which only few variants have been characterized and no standard molecules are available so far. Alternatively, they may correspond to completely new family of metabolites, which existence have been suggested by the observation of orphan NRPS/PKS clusters within various *Microcystis* genomes (Humbert *et al.*, 2013).

## 3. Discussion

### 3.1. Phylogeny and biogeography of Microcystis strains

The classification of species in genus *Microcystis* is still under large revision. Traditional morphological criteria including colony form, mucilage structure, cell diameter, the density and organization of cells within the colony, pigment content and life cycles are used for morphospecies recognition (Komarek 2016). Five dominant morphospecies (*M. aeruginosa, M. ichthyoblabe*, *M. viridis*, *M. novacekii*, *M. wesenbergii*) of the genus *Microcystis* were suggested by Ostuka *et al.* (2001). However, with the development of molecular and biochemical markers, some contradictory results of *Microcystis* taxonomy have been found. For instance, 16S rRNA analysis revealed no differences among the morpho-species (Otsuka *et al.*, 2000). In consideration of both morphological and molecular markers, it has been suggested to classify *Microcystis* into three groups: the small cell-size group composed of *M. ichthyoblabe* and *M. flos-aquae*, the middle cell-size group based on *M. aeruginosa* (incl. *M. novacekii*) and the large cell-size group represented by *M. wesenbergii* (Whitton, 2012). According to your phylogenetic reconstruction based on 16S-16S/23S ITS fragment that is in accordance with previous observation (Otsuka *et al.*, 1999), the studied *Microcystis* strains were roughly divided into two groups; the *M. wesenbergii/viridis* group and the *M. aeruginosa* group, in relation with the relative size of the colonies observed under microscopes. In addition, the *mcyA*+ and *mcyA*-strains appears broadly disperse on both group, unspecifically. These observations confirm that the phylogenetic relationship between different strains within the genus of *Microcystis* do not correspond to the possession of *mcy* (Tillett et al., 2001). Therefore, the toxic potential, through the MC synthesis, of each strain should assess regardless of its phylogenetical position.

### 3.2. *Biogeography and genetic diversity of* Microcystis *strains*

Despite the very high similarity of *Microcystis* 16S rRNA sequences (>99.5%), low synteny and large genomic heterogeneity have been retrieved from the investigation of *Microcystis* genomes, deciphering a large cryptic diversity from various strains collected from different sites and continents (Humbert *et al.*, 2013). At an other large geographic scale that in our analysis, genetic comparison of various *Microcystis* strains isolated from different Asian and European lakes based on 16S-ITS fragments does not show local-specific clustering effect indicating that the genetic distance between different genotypes from the same lake can be greater than between strains from very distanced environments (Humbert *et al.*, 2005; Haande *et al.*, 2007). Our data on the molecular heterogeneity observed by metabolic fingerprinting between some strains originating from the same site (for example between PMC 810.12 and PMC 816.12, Champs-sur-Marne, France) also support this hypothesis. The fact that some *Microcystis* genotypes seem to be distributed worldwide was also previously reported for bacterioplankton species (Zwart *et al.*, 1998), suggesting that these organisms may possess peculiar physiological capabilities and/or large dispersal capabilities enabling them to compete successfully in a wide range of freshwater environments.

### 3.3. *MC-producing* versus *non-MC producing metabolic pattern of* Microcystis *strains*

The metabolome diversity has been also used as molecular characters that can help at the discrimination of various chemotypes (Ivanizevic *et al.*, 2011), in addition to classical genotyping approach that sometimes lack of reliable characters for phylogenetic relationship discrimination. Few analytical methods have been experimented for chemotaxonomic characterization of cyanobacteria, based for example on fatty acid compositions (Gugger *et al.*, 2002), or more recently on ribosomal protein analysed globally by MALDI-TOF in *Microcystis aeruginosa* (Sun *et al.*, 2016). Interestingly, this latter chemotaxonomic approach was able to group all the MC-producing strains in two distinct clades when the non MC-producing strains were segregate in three other distinct clades. In a previous work, Martins and collaborators (2009) have analysed the metabolite diversity of various *Microcytis aeruginosa* strains originated from Portuguese water supplies using MALDI-TOF MS, and were able to observe a MC production in almost half of the strains investigated. These observations also illustrate the fact that MC-producing clones could subsist in various environments, despite the important energetic cost required for MC gene cluster replication and the translation of its mega-enzyme complex (Briand *et al.*, 2012). The biological advantage of producing MCs for some clones still remains an enigma, as the functional role of MC remaining uncharacterized (Agha and Quesada 2014; Gan *et al.*, 2012).

In a similar manner our molecular fingerprint approach based on global metabolome profiling using ESI-Qq-TOF discriminate clearly MC-producing strains from others. Taken together, these observations suggest that shotgun mass spectrometry chemotyping of cyanobacteria could constitute a promising tool for the characterization of rapid biomarkers aim at the toxicological assessment of strains isolated from the field. The MC production could constitute a singular treat, that could constitute on of the key drivers of the global metabolic diversity of *Microcystis* strains, suggesting that MC could play a keystone function in cyanobacterial metabolite production. Indeed, it has previously hypothesized according to metabolomic observation that the character of MCs production could be compensate in strains not producing MCs by the production of other metabolites, such as aeruginosamines (Martins et al., 2009) for unknown biological reasons (Briand *et al.*, 2016a; Tonk *et al.*, 2009). Such secondary metabolic compensatory mechanisms, between and within peptide classes, were previously suspected for *Microcystis* (Martins *et al.*, 2009) or *Planktothrix* (Tonk *et al.*, 2005) in response to various growth conditions. However, further investigations with wider sampling are now required in order to increase the data set that could better help to test such metabolite functional hypothesis.

### 3.3. Secondary metabolite diversity within known metabolite families

The molecular cluster identified by GNPS approach can be annotated thank to match with spectral databases or the presence of standard molecules present in the request as additional samples analysed similarly (Yang *et al.*, 2013). In our hands, the global molecular networks obtained from the 24 strains MS/MS dataset present various cluster that could have been annotated accordingly as corresponding to main cyanobacterial metabolite families. We assumed that the nodes of these clusters which molecular masses is similar to already known cyanobacterial metabolite very likely correspond to these specific metabolites or alternatively to isobaric analogues from the same family. In addition, all other nodes from those clusters that do not correspond to neither standard, nor known analogues, could be considered as new analogues that may correspond to new putative variants that still have to be investigated. These observations are in accordance with previous investigation indicating that different *Microcystis* strains can produce such various known and unknown secondary metabolites, according to both genetic or targeted metabolome analyses (Humbert *et al.*, 2013; Martins *et al.*, 2009; Welker *et al.*, 2004; Welker *et al.*, 2006). Surprisingly, rare metabolite family such as aeruginosamine are not successfully detected in our analysis. Indeed, only few molecules belonging to this family have been yet described (supp. Table 1) and their MS/MS fragmentation pattern has not been deeply characterized. In addition, the lack of available standard molecule and of knowledge on their respective fragmentation patterns makes aeruginosamines challenging to be simply annotate with our GNPS approach.

However, our analyses reveal the large molecular diversity of *Microcystis* metabolites, according to the various new variants of both cyanobacterial metabolite families that remain to be characterized. The observation of various uncharacterized cluster also suggest that new metabolite families are needed to be discovered and described from this taxa, and that further effort are still required. So far, *Microcystis* represent one of the most studied genera for its production of various metabolite families. However, the biological functions play by these molecules remains enigmatic and their overgrowing molecular diversity revealed by global approach, such as GNPS global metabolomic investigation, constitutes one of the questioning paradox in the field of evolution and diversity microbiology.

### 3.4. Unknown metabolite families

Although, the strains used in this study were not cultured in stringent axenic conditions, no noticeable contamination by fungi or heterotroph bacteria could have been detected during the systematic screening of all strains under light microscope prior to the experiment. In addition, a previous metabolome analyses in PCC 7806 grown under axenic or non-axenic condition do not detected any variation in the metabolite produced by the cyanobacteria (Briand *et al.*, 2016b). We assume that the metabolite profiles observed here for the 24 strains are characteristic of the cyanobacteria them self and that the different metabolite observed in this study, comprising the unknown metabolite clusters highlighted by the network analysis, are genuine produced by the cyanobacteria themselves.

The non-annotated cluster observed in our GNPS analysis can potentially correspond to novel variant of known cyanotoxin or to completely new family of cyanobacteria metabolite. Indeed, Humbert and co-workers (2013) have shown that the genome of ten *Microcytis* strains exhibits at least three orphan clusters with specific NRPS/PKS signature that are virtually synthesizing so far undescribed metabolite family. One could speculate that the unknown clusters observed with GNPS approach can correspond to such novel metabolite family, and structural elucidation of an expending number of novel metabolites revealed by molecular networking are currently been performed on various cyanobacteria (Boudreau *et al.*, 2015).

## 4. Conclusions

Innovative approaches based on shotgun metabolomic analyses using high resolution mass spectrometry, as those performs in this study, seems to provide a large panel of information on cyanobacteria chemical diversity relevant for evolutive, ecological and toxicological purposes, and represents an interesting and relatively easy-to-perform alternative to genome sequencing for metabolite and/or toxic potential descriptions of cyanobacterial strains.

Global molecular network also allows to depict the chemical diversity of the *Microcystis* metabolome in an interesting manner, with comparison with classical natural product chemistry approaches described so far, as in our hand above half of the analytes described in the global molecular network seems to correspond to metabolites belonging to potentially new variants of known families or even to family members (presenting original fragmentation patterns) that are still to be described at the structural and toxicological/bioactivity levels.

## 5. Material and methods

### 6.1. *Sampling, isolation and cultivation of* Microcystis *monoclonal strains*

The study was carried out with 24 monoclonal non-axenic cultures of *Microcystis* spp. maintained at 25°C in 15-mL vessels with Z8 media in the PMC (Paris Museum Collection) of living cyanobacteria (http://www.mnhn.fr/fr/collections/ensembles-collections/ressources-biologiques-cellules-vivantes-cryoconservees/microalgues-cyanobacteries). Larger volume of all strains was simultaneously cultivated during one month in triplicates in 50 mL Erlenmeyer’s vessels at 25°C using a Z8 medium with a 16 h: 8 h light/dark cycle (60 μmol.m-2.s-1). All trains were investigated for their MC production by Adda-microcystin AD4G2 ELISA kit (Abraxis). Cyanobacterial cells were centrifuged (at 4,000 g for 10 min), then freeze-dried and weighted, and stored at −80°C prior to DNA and metabolite analyses.

### 6.2. DNA-extraction, PCR, sequencing and Phylogenetic analyses

DNA was extracted with Qiagen Kit (Cat N°69506) according to manufacturer’s instructions. Presence and condition of the extracted DNA was confirmed by observing the 260/280-nm ratio and the absorbance spectra between 200 and 800 nm using a nanodrop spectrophotometer (Safas, Monaco). PCR reaction was performed with *mcyA* specific primers developed for *Microcystis* (*mcyA*_S AAAAACCCGCGCCCTTTTAC and *mcyA*_AS AGGCAGTTGGAGAATCACGG) in order to investigate the presence of this gene in the different strains. In parallel, the region containing a fragment 16S rRNA end of the 16S-23S ITS was amplified using primer couples previously described in Gugger and Hoffman (2004) and Iteman *et al.* (2000), respectively. The amplification was done in a reaction mixture containing 0.1 μL (100 μM) of each primer, 12.5 μL MyTaq Red Mix polymerase (Bioline®) and 2 μL (~200 ng) DNA sample. Final volume of the reaction mixture was 25 μL. The PCR product was sequenced (Genoscreen, France) using the same primers. The partial 16S-ITS sequences (above 1980-bp long) of all strains were deposited to GenBank (accession numbers xxxxx-xxxx).

The *Microcystis* 16S-23S ITS gene sequences were compared to a selection of similar (>93% identity) sequences retrieved from NCBI based on a standard nucleotide BLAST search (basic local alignment search tool). The sequences were aligned with CodonCode Aligner and non-homologous regions of the sequence alignment were manually deleted in BioEdit (Version 7.2.5). The phylogeny of the edited, aligned 16S-23S ITS sequences was performed in the program MEGA (Version 6). The trees based on maximum likelihood were constructed with 1000 bootstrap replicates, with the branch lengths iterated and global rearrangements done.

### 6.3. Metabolome biomass extraction and analysis by mass spectrometry

The 20 mL of biomasses of the 24 *Microcystis* strain cultures were centrifuged (4,000 rpm, 10 min), the culture media discarded, and then freeze-dried. The lyophilized cells were weighted then sonicated 2 min in acetonitrile/methanol/water (40/40/20) acidified at 0.1% of formic acid with a constant ratio of 100 μL of solvent for 1 mg of dried biomass, centrifuged at 4°C (12,000 g; 5 min). Two μL of the supernatant were then analyzed on an UHPLC (Ultimate 3000, ThermoFisher Scientific) coupled with a mass spectrometer (ESI-Qq-TOF Maxis II ETD, Bruker).

Ultra high performance liquid chromatography (UHPLC) was performed on 2 μL of each of the metabolite extracts using a Polar Advances II 2.5 pore C18 column (Thermo) at a 300 μL.min-1 flow rate with a linear gradient of acetonitrile in 0.1% formic acid (5 to 90% in 21 min). The metabolite contents were analyzed in triplicate for each strain using an electrospray ionization hybrid quadrupole time-of-flight (ESI-QqTOF) high resolution mass spectrometer (Maxis II ETD, Bruker) on positive simple MS or on positive autoMSMS mode with information dependent acquisition (IDA), on the 50-1500 *m/z* rang at 2 Hz or between 2-16 Hz speed, for MS and MS/MS respectively, according to relative intensity of parent ions, in consecutive cycle times of 2.5 s, with an active exclusion of previously analysed parents. The data were analyzed with the DataAnalysis 4.4 and MetaboScape 3.0 software for internal recalibration (<0.5 ppm), molecular feature search and MGF export. Peak lists were generated from MS/MS spectra between 1 and 15 min, with a filtering noise threshold at 0.1% maximal intensity and combining various charge states and related isotopic forms. Metabolite annotation was attempted according to the precise mass of the molecules and their respective MS/MS fragmentation patterns with regards to an in-house database of above 700 cyanobacteria metabolites and confirmed with few commercially available standard molecules analysed similarly in our platform.

### 6.4. Data and statistical analysis

Heatmap representation of the global metabolome of the 24 *Microcystis* spp. monoclonal strains was performed with Gene-E tool (https://software.broadinstitute.org/GENE-E/) using the relative quantification (pic area) of 2051 molecular features analysed on HR ESI-Qq-TOF using MetaboScape 3.0 (Bruker) with a >5000 counts and 400-2000 Da threshold, considering peak presents in at least 3 different trains and in at least 6 consecutive MS scans. Then, the hierarchical clustering was performed according to Bray-Curtis distance method. NMDS and PERMANOVA analyses were performed using MicrobiomeAnalyst platform (http://www.microbiomeanalyst.ca/) in order to investigate the influence of the species, the sampling localities and of the production of MCs, described as the variables, on the global metabolite distribution of the global metabolome observed on ESI-Qq-TOF for the 24 strains. Using the whole MSMS data (converted in mgf format) obtained for the 24 strains taken together, a molecular network was created using the online workflow at Global Natural Products Social molecular networking (GNPS) (http://gnps.ucsd.edu) (Wang *et al.*, 2016). The data were then clustered with MS-Cluster with a parent mass tolerance of 1.0 Da and an MS/MS fragment ion tolerance of 0.5 Da to create consensus spectra. Consensus spectra that contained less than one spectrum were discarded. A network was then created where edges were filtered to have a cosine score above 0.6 and more than five matched peaks. Further edges between two nodes were kept in the network only if each of the nodes appeared in each other’s respective top 10 most similar nodes. The spectra in the network were then searched against the GNPS spectral libraries. All matches kept between network spectra and library spectra were required to have a score above 0.6 and at least five matched peaks. The clustered spectra of the network were annotated by comparing monoisotopic mass to our in-house cyanobacteria metabolite databases according to MS and MS/MS fragmentation pattern matches. Molecular networks were visualized using Cytoscape 3.2.1.

## 6. Acknowledgements

This work was supported by grants from the CNRS Défi ENVIROMICS “Toxcyfish” project and from the ATM “Les micro-organismes, acteurs clef des écosystèmes” of the MNHN to Dr. Benjamin Marie. We would like to thank the French minister for the research for their financial supports Séverine Le Manach. The MS spectra were acquired at the Plateau technique de Spectrométrie de Masse Bio-organique, UMR 7245 Molécules de Communication et Adaptation des Micro-organismes, Muséum National d’Histoire Naturelle, Paris, France.

## Author contributions

SLM, ME, AC, CB and BM conceived and designed the experiments; CD isolated all new strains of the PMC; CD, SLM, AM, CDJ performed the analysis; SLM, CD, AM, SZ and BM treated the data. All authors wrote and reviewed the manuscript.

## Supplementary figures

**supplementary figure 2**: Representation of the analytes from the 24 *Microcystis* strains analysed by MS simple and MS/MS mode, exhibiting the good representativeness of analytes selected for MS/MS analyses. All analysed ions are represented according to their respective RT and *m/z* ratio. For MS/MS data, the size of the circle being representative of their maximum peak intensity.

**supplementary figure 2**: NMDS analysis of global metabolite patterns of the 24 *Microcystis* spp. monoclonal strains analysed using HR ESI-TOF, with PERMANOVA analyses performed on MicrobiomeAnalyst platform with Bray-Curtis index according to the MC production (left) and to the genera (right). “MC production”, “species”, and “locality” factor present significant impact on the global metabolome.

**Supplementary Figure 3**: Unknown cluster “1” highlighted by the GNPS analysis based on the MS/MS CID fragmentation spectra obtained from the 24 *Microcystis* strains. This cluster of uncharacterized molecules that present high fragmentation similarity main correspond to a new kind of metabolites that still need to be characterized. Analytes detected in *M. aeruginosa* or *M. wessenbergii/viridis* strains only are indicated in green and blue, respectively, when analytes detected in both genera are in orange.

**Supplementary Figure 4**: Unknown cluster “2” highlighted by the GNPS analysis based on the MS/MS CID fragmentation spectra obtained from the 24 *Microcystis* strains. This cluster of uncharacterized molecules that present high fragmentation similarity main correspond to a new kind of metabolites that still need to be characterized. Analytes detected in *M. aruginosa* and *M. wessenbergii/viridis* strains are indicated in orange.

**Supplementary Figure 5**: Unknown cluster “3-5” highlighted by the GNPS analysis based on the MS/MS CID fragmentation spectra obtained from the 24 *Microcystis* strains. This cluster of uncharacterized molecules that present high fragmentation similarity main correspond to a new kind of metabolites that still need to be characterized. Analytes detected in *M. wessenbergii/viridis* strains only are indicated in blue.

## References

Agha, R., Quesada, A. (2014) Oligopeptides as biomarkers of cyanobacterial subpopulations. Toward an understanding of their biolgical role. Toxins 6, 1929–50.

Boudreau, P., Monroe, E., Mehrotra, S., Desfor, S., Korabeynikov, A., Sherman, D., Murray, T., Gerwick, L., Dorrestein, P., Gerwick, W. (2015) Marine cyanobacterium Moorea producens JHB through orthogonal natural products workflows. PLoS ONE 10:e0133297.

Briand, E., Escoffier, N., Straub, C., Sabart, M., Quiblier, C., Umbert, J-F. (2009) Spatiotemporal changes in the genetic diversity of a bloom-forming Microcystis aeruginosa (cyanobacteria) population. ISME J. 3, 419–429.

Briand, E., Bomans, M., Quiblier, C., Saleçon, M-J., Humbert, J-F. (2012) Evidence of the cost of the production of Microcystins by Microcystis aeruginosa under different light and nitrate environmental conditions. PLoS ONE 7:e29981.

Briand, E., Bormans, M., Gugger, M., Dorrestein, PC., Gerwick, W. (2016a) Changes in secondary metabolic profiles of Microcystis aeruginosa strains in response to intraspecific interactions. Environmental Microbiology 18:384–400.

Briand, E., Humbert, J-F., Tambosco, K., Bormans, M., Gerwick, W; (2016b) Role of bacteria in the prodcution and degradation of Microcystis cyanopeptides. Microbiology Open 3:343.

Carey, C.C., Ibelings, B.W., Hoffmann, E.P., Hamilton, D.P., Brookes, J.D. (2012) Eco-physiological adaptations that favour freshwater cyanobacteria in a changing climate. Water Res. 46, 1394–1407. doi:10.1016/j.watres.2011.12.016

Carmichael, W. (2008) Cyanobacterial Harmful Algal Blooms: State of the Science and Research Needs. Adv. Exp. Med. Biol. 619, 831–53. doi:10.1007/978-0-387-75865-7

Catherine, A., Bernard, C., Spoof, L., Bruno, M. (2017). Microcystins and Nodularins. In Handbook of Cyanobacterial Monitoring and Cyanotoxin Analysis (eds J. Meriluoto, L. Spoof and G. A. Codd). doi:10.1002/9781119068761.ch11

Codd, G.A., Morrison, L.F., Metcalf, J.S. (2005) Cyanobacterial toxins: risk management for health protection. Toxicol. Appl. Pharmacol. 203, 264–72. doi:10.1016/j.taap.2004.02.01

Dittmann, E., Gugger, M., Sivonen, K., Fewer, D.P. (2015) Natural Product Biosynthetic Diversity and Comparative Genomics of the Cyanobacteria. Trends Microbiol. 23, 642–652. doi:10.1016/j.tim.2015.07.008

Gan, N., Xiao, Y., Zhu, L., Wu, Z., Liu, J., Hu, C., Song, L., 2012. The role of microcystins in maintaining colonies of bloom-forming Microcystis spp. Environ. Microbiol. 14, 730–742. doi:10.1111/j.1462-2920.2011.02624.x

Gugger, M., Hoffmann, L. (2004) Polyphyly of true branching cyanobacteria (Stigonematales). Int J Syst Evol Microbiol 54, 349–57.

Gugger, M., Lyra, C., Suominen, I., Tsitko, I., Humbert, J-F., Salkinoja-Salonen, M., Sivonen, K. (2002) Cellular fatty acids as chemotaxonomic makers of the genera Anabaena, Aphanizomenon, Microcystis, Nostoc and Planktothrix. Int. J. System. Evol. Microbiol. 52, 1007–1015.

Guljamow, A., Kreische, M., Ishida, K., Liaimer, A., Altermark, B., Bähr, L., Hertweck, C., Ehwald, R., Dittmann, E. (2017) High-density cultivation of terrestrial Nostoc strains leads to reprogramming of secondary metabolome. Appl Environ Microbiol. 83 :e01510–17.

Haande, S., Ballot, A., Rohrlack, T., Fastner, J., Wiedner, C., Edvardsen, B. (2007) Diversity of Microcystis aeruginosa isolates (Chroococcales, Cyanobacteria) from East-African water bodies. Arch. Microbiol. 188, 15–25. doi:10.1007/s00203-007-0219-8

Harke, M.J., Steffen, M.M., Gobler, C.J., Otten, T.G., Wilhelm, S.W., Wood, S.A., Paerl, H.W. (2016) A review of the global ecology, genomics, and biogeography of the toxic cyanobacterium, Microcystis spp. Harmful Algae 54, 4–20. doi:10.1016/j.hal.2015.12.007

Harke, M.J., Jankowiak, J.G., Morrell, B.K., Gobler, C.J. (2017) Transcriptomic Responses in the Bloom-Forming Cyanobacterium Microcystis Induced during Exposure to Zooplankton. Appl Environ Microbiol. 15;83(5).

Holland, A., Kinnear, S. (2013) Interpreting the possible ecological role(s) of cyanotoxins: compounds for competitive advantage and/or physiological aide? Mar. Drugs 11, 2239–58. doi:10.3390/md11072239

Humbert, J-F., Duris-Latour, D., Le Berre, B., Giraudet, H., Salençon, M.J. (2005) Genetic diversity in Microcystis populations of a French storage reservoir assessed by sequencing of 16S-23S rRNA intergenic spacer. Microbiol Ecology 49, 308–314.

Humbert, J-F., Barbe, V., Latifi, A., Gugger, M., Camteau, A., Coursin, T., Lajus, A., Castelli, V., Oztas, S., Samson, G., Longin, C., Medigue, C., Tandeau de Marsac, N. (2013) A tribute to disorder in the genome of the bloom-forming freshwater cyanobacterium Microcystis aeruginosa. PLoS ONE 8: e70747.

Iteman, I., Rippka, R., Tandeau de Marsac, N., et al. (2000) Comparison of conserved structural and regulatory domains within divergent 16SrRNA–23S rRNA spacer sequences of cyanobacteria. Microbiology 146:1275–86.

Ivanisevic, J., Thomas, O., Lejeune, C., Chavaldonné, P., Perez, T. (2011) Metabolic fingerprinting as an indicator of biodiversity: towards understanding inter-specific relationships among Homoscleromorpha sponges. Metabolomics 7: 289–304.

Komárek, J. (2016). A polyphasic approach for the taxonomy of cyanobacteria: principles and applications. European Journal of Phycology, 51(3), 346–353.

Liu, Y., Xu, Y., Wang, Z., Xiao, P., Yu, G., Wang, G., Li, R. (2016) Dominance and succession of Microcystis genotypes and morphotypes in Lake Taihu, a large and shallow freshwater lake in China. Environ. Pollut. 219, 399–408. doi:10.1016/j.envpol.2016.05.021

Ma, J., Qin, B., Paerl, H.W., Brookes, J.D., Hall, N.S., Shi, K., Zhou, Y., Guo, J., Li, Z., Xu, H., Wu, T., Long, S. (2016) The persistence of cyanobacterial (Microcystis spp.) blooms throughout winter in Lake Taihu, China. Limnol. Oceanogr. 61, 711–722. doi:10.1002/lno.10246

Martins, J., Saker, M.L., Moreira, C., Welker, M., Fastner, J., Vasconcelos, V.M. (2009) Peptide diversity in strains of the cyanobacterium Microcystis aeruginosa isolated from Portuguese water supplies. Appl. Microbiol. Biotechnol. 82, 951–961.

Martins, J., Vasconcelos, V. (2015) Cyanobactins from cyanobacteria: Current genetic and chemical state of knowledge. Mar. Drugs 13, 6910–6946. doi:10.3390/md13116910

Otsuka, S., Suda, S., Shibata, S., Oyaizu, H., Matsumoto, S., Watanabe, M.M (2001) A proposal for the unification of five species of the cyanobacterial genus Microcystis Kützing ex Lemmermann 1907 under the rules of the Bacteriological Code. Int J Syst Evol Microbiol. 51(Pt 3), 873–9.

Otsuka, S., Suda, S., Li, R., Matsumoto, S., Watanabe, M.M. (2000) Morphological variability of colonies of Microcystis morphospecies in culture. J Gen Appl Microbiol. 46(1), 39–50.

Otsuka, S., Suda, S., Li, R., Watanabe, M. (1999) Phylogenetic relationships between toxic and non-toxic strains of the genus Microcystis based on 16S to 23S internal transcribed spacer sequence, FEMS Microbiol. Lett. 172 (1),15–21.

Paerl, HW. (2018) Mitigating Toxic Planktonic Cyanobacterial Blooms in Aquatic Ecosystems Facing Increasing Anthropogenic and Climatic Pressures. Toxins 8;10(2).

Paerl, H.W., Hall, N.S., Calandrino, E.S. (2011) Controlling harmful cyanobacterial blooms in a world experiencing anthropogenic and climatic-induced change. Sci. Total Environ. 409, 1739–45. doi:10.1016/j.scitotenv.2011.02.001

Pearson, L., Mihali, T., Moffitt, M., Kellmann, R., Neilan, B. (2010) On the chemistry, toxicology and genetics of the cyanobacterial toxins, microcystin, nodularin, saxitoxin and cylindrospermopsin. Mar. Drugs 8, 1650–80. doi:10.3390/md8051650

Šejnohová, L., Maršálek, B. (2012) Microcystis, in: Ecology of Cyanobacteria II: Their Diversity in Space and Time. Springer Netherlands, Dordrecht, p. 195–228. doi:10.1007/978-94-007-3855-3_7

Shih, P.M., Wu, D., Latifi, A., Axen, S.D., Fewer, D.P., Talla, E., Calteau, A., Cai, F., Tandeau de Marsac, N., Rippka, R., Herdman, M., Sivonen, K., Coursin, T., Laurent, T., Goodwin, L., Nolan, M., Davenport, K.W., Han, C.S., Rubin, E.M., Eisen, J. a, Woyke, T., Gugger, M., Kerfeld, C. (2013) Improving the coverage of the cyanobacterial phylum using diversity-driven genome sequencing. Proc. Natl. Acad. Sci. U. S. A. 110, 1053–8. doi:10.1073/pnas.1217107110

Sivonen, K., Leikoski, N., Fewer, D.P., Jokela, J. (2010) Cyanobactins-ribosomal cyclic peptides produced by cyanobacteria. Appl. Microbiol. Biotechnol. 86, 1213–1225. doi:10.1007/s00253-010-2482-x

Sukenik, A., Quesada, A., Salmaso, N. (2015) Global expansion of toxic and non-toxic cyanobacteria: effect on ecosystem functioning. Biodivers. Conserv. 4:889–908.

Sun, L-W., Jiang, W-J., Sato, H., Kawachi, M., Lu, X-W. (2016) Rapid classification and identification of Microcystis strains using MALDI-TOF MS and polyphasic analysis. PLoS ONE 11: e0156275.

Tillett, D., Parker, D. L., & Neilan, B. A. (2001). Detection of toxigenicity by a probe for the microcystin synthetase A gene (mcyA) of the cyanobacterial genus Microcystis: comparison of toxicities with 16S rRNA and phycocyanin operon (phycocyanin intergenic spacer) phylogenies. Applied and environmental microbiology, 67(6), 2810–2818.

Tonk, L., Visser, P.M., Christiansen, G., Dittmann, E., Snelder, E.O.F.M., Wiedner, C., Mur, L.R., Huisman, J. (2005) The microcystin composition of the cyanobacter-ium Planktothrix agardhii changes towards a more toxic variant with increasing light intensity. Appl. Environ. Microbiol. 71, 5177–5181.

Tonk, L., Welker, M., Huisman, J., Visser, P.M. (2009) Production of cyanopeptolins, anabaenopeptins, and microcystins by the harmful cyanobacteria Anabaena 90 and Microcystis PCC 7806. Harmful Algae 8, 219–224. doi:10.1016/j.hal.2008.05.005

Via-Ordorika, L., Fastner, J., Kurmayer, R., Hisbergues, M., Dittmann, E., Komarek, J., Erhard, M., Chorus, I. (2004) Distribution of microcystin-producing and non-microcystin-producing Microcystis sp. in European freshwater bodies: detection of microcystins and microcystin genes in individual colonies. Syst Appl Microbiol. 27(5), 592–602.

Wang, H., Fewer, D.P., Holm, L., Rouhiainen, L., Sivonen, K. (2014) Atlas of nonribosomal peptide and polyketide biosynthetic pathways reveals common occurrence of nonmodular enzymes. Proc Natl Acad Sci U S A. 111(25), 9259–64.

Welker, M., Brunke, M., Preussel, K., Lippert, I., von Döhren, H. (2004) Diversity and distribution of Microcystis (cyanobacteria) oligopeptide chemotypes from natural communities studies by single-colony mass spectrometry. Microbiology 150, 1785–1796. doi:10.1099/mic.0.26947-0

Welker, M., Von Döhren, H. (2006) Cyanobacterial peptides - Nature’s own combinatorial biosynthesis. FEMS Microbiol. Rev. 30, 530–563. doi:10.1111/j.1574-6976.2006.00022.x

Welker, M., Maršálek, B., Šejnohová, L., von Döhren, H. (2006) Detection and identification of oligopeptides in Microcystis (cyanobacteria) colonies: Toward an understanding of metabolic diversity. Peptides 27, 2090–2103. doi:10.1016/j.peptides.2006.03.014

Welker, M., Ejnohová, L., Némethová, D., von Döhren, H., Jarkovsky, J., Marsálek, B.(2007) Seasonal shifts in chemotype composition of Microcystis sp. communities in the pelagial and the sediment of a shallow reservoir. Limnol. Oceanogr. 52, 609–619. doi:10.4319/lo.2007.52.2.0609

Welker, M., Dittmann, E., Von Döhren, H. (2012) Cyanobacteria as a source of natural products. Methods Enzymol. 517, 23–46. doi:10.1016/B978-0-12-404634-4.00002-4

Whitton, B.A. (2012) Ecology of Cyanobacteria II, Ecology of Cyanobacteria II: Their Diversity in Space and Time. Springer Netherlands, Dordrecht. doi:10.1007/978-94-007-3855-3

Yang, J.Y., Sanchez, L.M., Rath, C.M., Liu, X., Boudreau, P.D., Bruns, N., Glukhov, E., Wodtke, A., De Felicio, R., Fenner, A., Wong, W.R., Linington, R.G., Zhang, L., Debonsi, H.M., Gerwick, W.H., Dorrestein, P.C. (2013) Molecular networking as a dereplication strategy. J. Nat. Prod. 76, 1686–1699. doi:10.1021/np400413s

Zak, A., Kosakowska, A. (2016) Cyanobacterial and microalgal bioactive compounds-the role of secondary metabolites in allelopathic interactions. Oceanol. Hydrobiol. Stud. 45, 131–143. doi:10.1515/ohs-2016-0013

Zurawell, R.W., Chen, H., Burke, J.M., Prepas, E.E. (2005) Hepatotoxic cyanobacteria: a review of the biological importance of microcystins in freshwater environments. J. Toxicol. Environ. Health. B. Crit. Rev. 8, 1–37. doi:10.1080/10937400590889412

Zwart, G., Hiorns, W.D., Methe, B.A., van Agterveld, M.P., Huismans, R., Nold, S.C., Zehr, J.P., Laanbroek, H.J. (1998) Nearly identical 16S rRNA sequences recovered from lakes in North America and Europe indicate the existence of clades of globally distributed freshwater bacteria. Syst Appl Microbiol. 21(4):546–556.

